# Dual Targeting of STING and PI3Kγ Eliminates Regulatory B Cells to Overcome STING Resistance for Pancreatic Cancer Immunotherapy

**DOI:** 10.1101/2024.02.14.580378

**Authors:** Chengyi Li, Shuai Mao, Hongyi Zhao, Miao He, Meilin Wang, Zhongwei Liu, Hanning Wen, Zhixin Yu, Bo Wen, Djibo Mahamadou, Jinsong Tao, Yingzi Bu, Wei Gao, Duxin Sun

## Abstract

The immune suppression in tumors and lymph nodes of pancreatic ductal adenocarcinoma (PDAC), regulated by suppressive myeloid cells and regulatory B (Breg) cells, hinders the effectiveness of immunotherapy. Although STING agonists activate myeloid cells to overcome immune suppression, it expands Breg cells, conferring STING resistance in PDAC. We discovered that blocking PI3Kγ during STING activation abolished IRF3 phosphorylation to eliminate Breg cells, while PI3Kγ inhibition sustained STING-induced IRF3 phosphorylation to preserve STING function in myeloid cells. Therefore, we developed a dual functional compound SH-273 and its albumin nanoformulation Nano-273, which stimulates STING to activate myeloid cells and inhibits PI3Kγ to eliminates Breg cells overcoming STING resistance. Nano-273 achieved systemic antitumor immunity through intravenous administration, which decreases Breg cells and remodels microenvironment in tumors and lymph nodes. Nano-273, combined with anti-PD-1, extended median survival to 200 days in transgenic KPC PDAC mice (KrasG12D-P53R172H-Cre), offering potential for PDAC treatment.

## Introduction

Pancreatic ductal adenocarcinoma (PDAC), the deadliest cancer type, has only an 12% five-year survival rate, underscoring a critical need for more effective treatments^1^. The only two available first-line therapeutic regimens for pancreatic cancer are FOLFIRINOX (leucovorin, fluorouracil, irinotecan, and oxaliplatin) and a combination of Abraxane (an albumin-based nanoparticle of paclitaxel) with gemcitabine. However, these treatments offer limited efficacy^2–4^. Notably, while immunotherapies targeting PD-1/PD-L1 have shown remarkable success in various other cancer types, their effectiveness in PDAC patients remains minimal^5,6^.

The poor efficacy of immunotherapy in PDAC is hindered by the immunosuppressive microenvironment in both tumors and lymph nodes, which consists both myeloid cells and regulatory B (Breg) cells. The myeloid cell-driven immune suppression in PDAC, such as tumor-associated macrophages (TAMs), myeloid-derived suppressor cells (MDSCs) and inactivate dendritic cells (DCs), is well-established^6,7^.Yet, the Breg cell-driven immune suppression in PDAC has only recently been recognized^8–11^. Recent studies demonstrate that PDAC has significantly high Breg cell populations, exacerbating immune suppression^8,9^ and contributing to negative patient outcomes during PDAC immunotherapy^8–10^.

However, current immunomodulators to counter immune suppression primarily focus on targeting myeloid cells^6,12^, without effectively regulating Breg cells^7,13^ in both tumors and lymph nodes. For instance, STING agonist, as one of the most effective immunomodulators, overcomes myeloid cell-driven immune suppression by stimulating type I interferon signaling^14^, but STING agonist simultaneously expands suppressive Breg cells^11^, which represents an intrinsic resistance mechanism in PDAC^13^. More importantly, the opposite effect of STING agonist in myeloid cells and in Breg cells depends on the same pathway through IRF3 phosphorylation^11^. In myeloid cells, STING activation increases IRF3 phosphorylation promoting type I interferon expression. This repolarizes TAMs into the M1 phenotype and stimulates dendritic cells (DCs) to activate cytotoxic CD8+ T cells^14^. However, in Breg cells, STING-induced IRF3 phosphorylation promotes IL-35 and IL-10 gene expression, which expands Breg cells for immune suppression^8,11,15^. Therefore, there is a pressing need to develop strategies to overcome the immune suppression by simultaneously activating myeloid cells and eliminating Breg cells in both tumors and lymph nodes in PDAC.

In this study, we discovered that blocking PI3Kγ during STING activation abolished IRF3 phosphorylation in B cells, thus decreasing Breg cells, while sustaining IRF3 phosphorylation in myeloid cells to preserve STING function. Based on the finding, we developed a dual functional compound SH-273 by stimulating STING and inhibiting PI3Kγ, which preserves STING function to activate myeloid cells and eliminates Breg cells to overcome STING resistance. To achieve systemic antitumor immunity, we prepared albumin nanoformulation of SH-273 (Nano-273) for intravenous administration to decrease Breg cells and remodel microenvironment in tumors and lymph nodes^11^, a goal not attainable through local intratumorally injection of STING agonists^13,14^. The in vivo efficacy of Nano-273 was evaluated in transgenic pancreatic cancer mouse model (KPC, KrasG12D, P53R172H, Pdx1-Cre) to extend median survival to 200 days, where no toxicity was observed at therapeutic dose regimen. Single-cell analysis and flow cytometry was used to evaluate Nano-273 to remodel microenvironment and eliminate Breg cells. These data suggest that Nano-273 is a promising candidate for pancreatic cancer immunotherapy, by eliminating Breg cells to overcome STING resistance and counteract suppressive microenvironment in both tumors and lymph nodes.

## Results

### STING agonists expand Breg cells in PDAC Mice

Recent studies suggest that immunosuppression in PDAC is mediated by both immune suppressive regulatory B cells and myeloid cells ^6,7,11–13^. To validate the presence of Breg cells in PDAC mouse model, we analyzed their frequency in tumors and lymph nodes of KPC (KrasG12D, P53R172H, Pdx1-Cre) transgenic mice, which spontaneously develop pancreatic adenocarcinoma in comparison with normal C57BL/6 mice. Our results revealed a significant increase in both IL35+ and IL10+ Breg cell frequencies in the tumor, lymph nodes of KPC mice compared to normal mice. Specifically, Breg cell frequencies were 4.6 and 36.2 times higher in KPC mouse tumors (10.2% and 19.9%) than in pancreatic tissues of normal mice (2.2% and 0.55%), 32.4 and 310 times higher in KPC mouse lymph nodes (22% and 18.6%) than in normal lymph nodes (0.68% and 0.06%). (**Fig. 1a, 1b**). Additionally, a high level of M2 macrophages was also observed in the tumor, lymph nodes, and spleen of PDAC mice (**Fig. S1**).

**Fig. 1.**
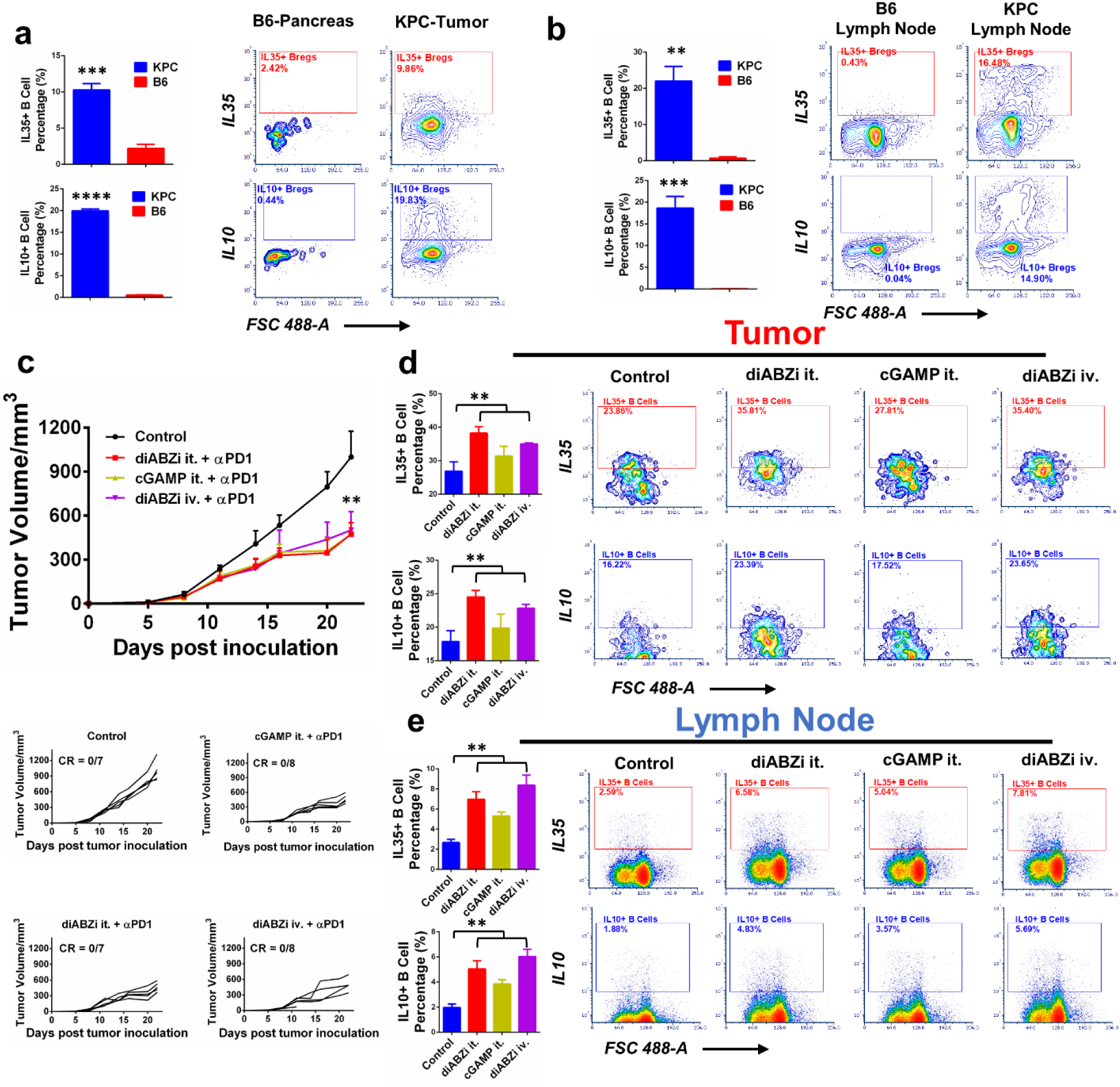
STING agonists expand Breg cells in PDAC Mice. Quantification and representative flow cytometry analysis of IL35^+^ and IL10^+^ Breg cells in KPC mice pancreatic tumor (**a**) and lymph node (**b**) in KPC mice and C57BL/6 mice. Data for quantification are shown as mean ± SD, n = 3. (**c**) Antitumor efficacy of STING agonists in C57BL/6 mice inoculated with pancreatic tumor (KPC 6422 cell line). Average tumor volumes changes after treatment with PBS (control), diABZi (intratumorally injection, 20 ug/mouse) plus anti-PD-1 antibody (100 µg), cGAMP (intratumorally injection 10 ug/mouse) plus anti-PD-1 antibody (100 µg), and diABZi (intravenous injection 1.5 mg/kg) plus anti-PD-1 antibody (100 µg). The data represents the mean ± SD; n = 5. Quantification and representative flow cytometry analysis of IL35^+^ and IL10^+^ Breg cells in tumor (**d**) and lymph node (**e**) from mice inoculated with KPC 6422 cell in Fig. 1c. Statistical comparisons are based on one-way ANOVA, followed by post hoc Tukey’s pairwise comparisons or by Student’s unpaired T-test. The asterisks denote statistical significance at the level of * p < 0.05, ** p < 0.01, *** p < 0.001, **** p < 0.0001. ANOVA, analysis of variance; SD, standard deviation. For (**c**), Statistical comparisons are compared between STING agonists groups with control group.

Recent studies also suggest that STING agonists further expand Breg cells, contributing to intrinsic STING resistance in PDAC^8,11,15^. To confirm these, we evaluated the in vivo effects of two STING agonists, cGAMP and diABZi, in C57BL/6 mice implanted with PDAC KPC 6422 cells. Following five doses of STING agonists administered either intratumorally (it) or intravenously (iv), in combination with anti-PD-1, we observed minimal tumor growth inhibition (**Fig. 1c**). Both intratumorally and intravenously administrations of STING agonists further expanded IL35^+^ and IL10^+^ Breg cells in tumor tissues and lymph nodes **(Fig. 1d and 1e)**. This persistence of high level Breg cells may explain the limited efficacy of STING agonists combined with anti-PD-1 in KPC mice, which underscores the importance of eliminating Breg cells to overcome STING resistance in PDAC.

### Blocking PI3Kγ abolished STING-induced IRF3 phosphorylation in B cells, while PI3Kγ inhibition sustained STING-induced IRF3 phosphorylation in myeloid cells

To understand the expressions of STING and PI3Kγ in different tissues and immune cells, we first used western blot to detect their protein levels. The data showed that STING is present in tissues including tumor and lymph nodes of KPC mice, normal pancreas and normal lymph nodes in C57BL/6 mice, as well as different immune cells including B cells, bone marrow derived macrophages (BMDM), bone marrow derived dendritic cells (BMDM) in both KPC mice and normal mice (**Fig. 2a, 2b**), In comparison, PI3Kγ is also expressed in tumor and lymph node, as well as different immune cells. (**Fig. 2a, 2b**).

**Fig. 2.**
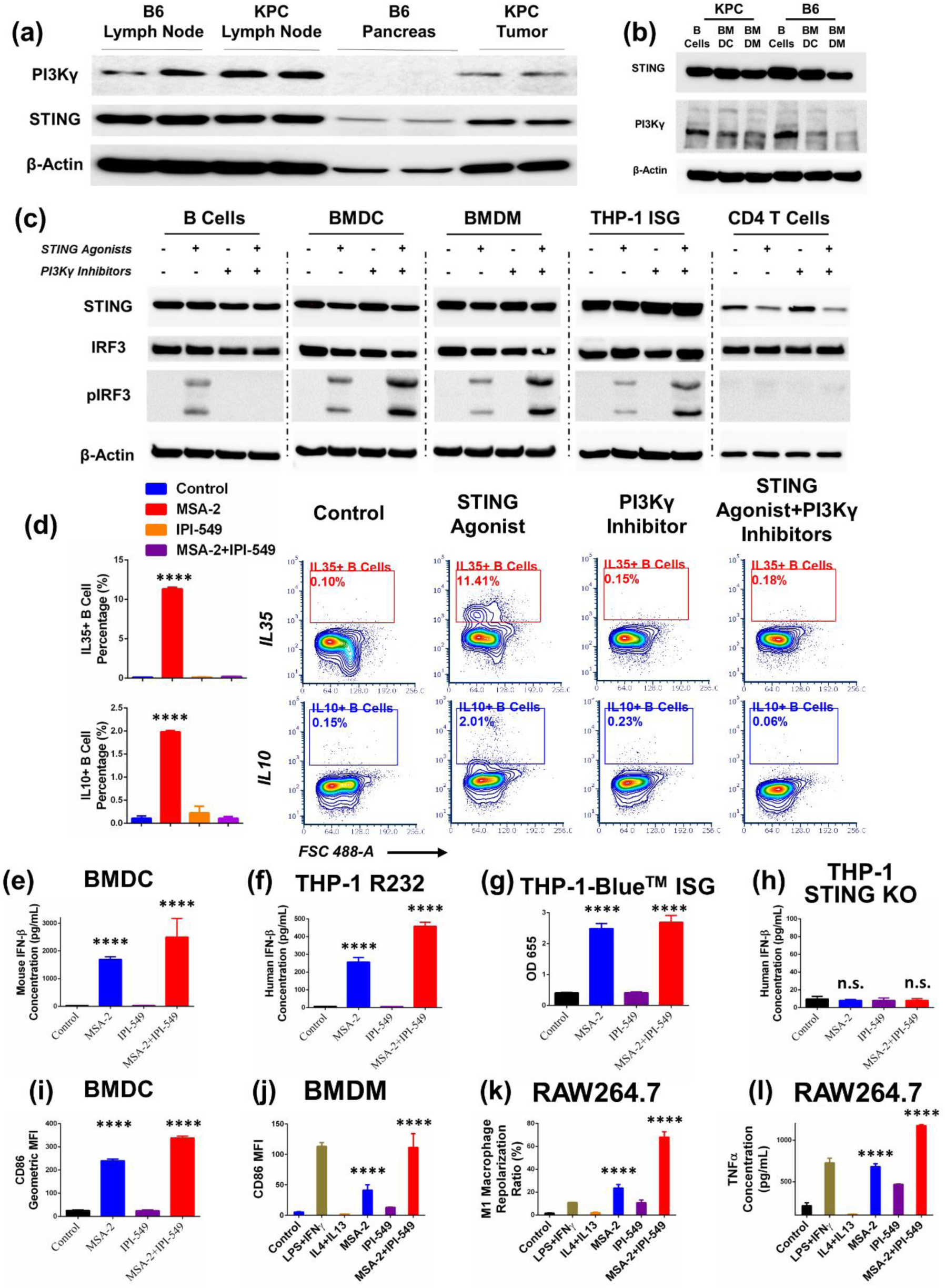
PI3Kγ inhibition abolished STING-induced IRF3 phosphorylation to eliminate STING-induced Breg cells expansion, while PI3Kγ inhibition sustained STING-induced IRF3 phosphorylation to preserve STING function in myeloid cells. (**a**) PI3Kγ and STING expression at lymph node and pancreatic tumor from KPC mice compared to C57BL/6 mice. (**b**) PI3Kγ and STING expression at B cell, bone marrow derived dendritic cells and bone marrow derived macrophages from KPC mice and C57BL/6 mice. (**c**) Phosphorylation of IRF3 at splenic B cells, bone marrow derived dendritic cells (BMDCs), bone marrow derived macrophages (BMDMs) and CD4 T cells derived from KPC mice and THP-1 hSTING^HAQ^ cells after treatments with or without MSA-2 or IPI-549. (**d**) Quantification and representative flow cytometry analysis of IL35^+^ and IL10^+^ Breg cells (from KPC spleen) after treatments with or without MSA-2 or IPI-549. (**e**) Quantification of mouse Interferon-β concentration of bone marrow derived dendritic cells (BMDCs) derived from C57BL/6 mice after treatments with or without MSA-2 or IPI-549. (**f, g**) Quantification of human Interferon-β concentration from THP-1 R232 hSTING^R232^ after treatments with or without MSA-2 or IPI-549. (**g**) Quantification of STING activation by reporter assay from THP1-Blue^TM^ ISG by measuring optical density at 655 nm after treatments with or without MSA-2 or IPI-549. (**h**) Quantification of human Interferon-β concentration from THP-1 hSTING^KO^ after treatments with or without MSA-2 or IPI-549. (**i, j**) Quantification of CD86 geometric mean fluorescent intensity of bone marrow derived dendritic cells (BMDCs) bone marrow derived macrophages (BMDMs) derived from C57BL/6 mice after treatments with or without MSA-2 or IPI-549. (**k**) Quantification of RAW264.7 M1 polarization ratios after treatments with or without MSA-2 or IPI-549. (**l**) Quantification of TNF-α concentration from RAW264.7 after treatments with or without MSA-2 or IPI-549. Data for quantification are shown as mean ± SD, n = 3. Statistical comparisons are based on one-way ANOVA, followed by post hoc Tukey’s pairwise comparisons or by Student’s unpaired T-test. The asterisks denote statistical significance at the level of **** p < 0.0001. ANOVA, analysis of variance; SD, standard deviation; n.s., no statistical significance. For (**d**) statistical comparisons are conducted between MSA-2 group with other groups. For (**e-l**), statistical comparisons are conducted between MSA-2 and MSA-2+IPI-549 groups with control group.

STING activation triggers IRF3 phosphorylation in myeloid cells, enhancing type I interferon expression, while in B cells, IRF3 phosphorylation boosts IL-10 and IL-35 expression, expands B regulatory (Breg) cells^8,11,15^. To investigate effect of PI3Kγ inhibition on STING-induced IRF3 phosphorylation across cell types, we used western blot to analyze IRF3 phosphorylation in the presence/absence of PI3Kγ inhibitor (IPI-549) and STING agonist (MSA-2) in B cells, Bone Marrow derived dendritic cell (BMDC) and bone marrow derived macrophage (BMDM) from KPC mice, and THP-1 hSTING^HAQ^ monocyte cell line, as well as CD4 T cells. STING activation by MSA-2 indeed increased IRF3 phosphorylation in B cells and myeloid cells (BMDC, BMDM, and THP-1 hSTING^HAQ^) (**Fig. 2c**), consistent with previous report^11^. Surprisingly, PI3Kγ inhibition by IPI-549 completely abolished STING-induced IRF3 phosphorylation in B cells, but it sustained STING-induced IRF3 phosphorylation in myeloid cells (**Fig. 2c**). These phenomena are not observed in CD4 T cells (**Fig. 2c**).

### Blocking PI3Kγ eliminated STING-induced Breg cells expansion, while PI3Kγ inhibition preserved STING induced myeloid cell activation

To further analyze how did PI3Kγ inhibition alter Breg cell populations, we used flow cytometry to monitor IL-35+ and IL-10+ Breg cells in the splenocytes isolated from KPC and STING knockout mice (Tmem173^-/-^), treated with or without the STING agonist MSA-2 and the PI3Kγ inhibitor IPI-549. The data showed that STING agonist (MSA-2) treatment significantly increased IL-35^+^ and IL-10^+^ Breg cells by 100 and 18-fold, but PI3Kγ inhibitor (IPI-549) completely eliminated these Breg cells (**Fig. 2d and S2**). To confirm this effect is STING-dependent, we also monitored IL-35+ and IL-10+ Breg cells in the splenocytes from STING knockout mice (Tmem173^-/-^ mice) under treatment of STING agonist (MSA-2) and PI3Kγ inhibitor (IPI-549). The data showed that neither MSA-2 nor IPI-549 has any effect on IL-35^+^ and IL-10^+^ Breg cells in STING knockout mice (**Fig. S3**), strongly suggesting STING dependence. Additionally, we found no significant induction of IL-35 and IL-10 in BMDCs, BMDMs, THP-1 cells, and CD4 T cells under similar treatment conditions (**Fig. S4-S8**), suggesting that STING-induced IL-35 and IL-10 expressions are exclusively in B cells.

To investigate whether PI3Kγ inhibition would impact STING activation in myeloid cells, we measured STING induced type I interferon expression in BMDC, monocyte THP1 hSTING^R232^, and THP-1-Blue^TM^ ISG (hSTING^HAQ^), and THP-1 hSTING^KO^ cells. The data shows that STING agonist MSA-2 stimulated IFN-beta secretion in these STING positive cells but not in STING knockout cells, while PI3Kγ inhibition (IPI-549) sustained STING activation in these STING positive cells (**Fig. 2e-2h**).

To study whether PI3Kγ inhibition would influence STING induced activation of myeloid cells, we measured dendritic cell activation and M1-macrophage polarization in BMDCs and BMDMs from KPC mice, as well as macrophage cell line RAW264.7 cells. The data showed that STING agonist (MSA-2) activated BMDCs (**Fig. 2i and S9**) and BMDMs (**Fig. 2j and S10**) as measured by CD86^+^ populations, stimulated M1-macrophage polarization (**Fig. 2k and S11**) and increased TNF-alpha secretion in macrophage RAW264.7 (**Fig. 2l**). PI3Kγ inhibition preserved the STING induced activation in these myeloid cells (**Fig 2e-2l**).

In summary, our results suggest that PI3Kγ inhibition abolished STING-induced IRF3 phosphorylation in B cells, thereby eliminating Breg cell expansion. However, PI3Kγ inhibition sustained STING-induced IRF3 phosphorylation in myeloid cells, promoting type I interferon production and facilitating the activation of myeloid cells.

### Dual functional compound SH-273 and its albumin nanoformulation inhibit PI3Kγ to eliminates Breg cells and stimulate STING function to activate myeloid cells

To simultaneously activate STING and inhibit PI3Kγ, we developed a dual functional compound (SH-273), and encapsulated it into albumin nanoformulation Nano-273 for intravenous administration to achieve systemic immunity **(Fig. 3a)**. The synthesis of SH-273 is described in **Scheme 1**. The dual functional compound SH-273 stimulated STING activity (IC50 29.6 nM) in THP-1 Blue ISG cells (STING reporter) (**Fig. S12**) and inhibited PI3Kγ (IC50 20.2 nM) (**Fig. S13**). SH-273 abolished the STING-induced IRF3 phosphorylation in B cells from splenocytes of KPC PDAC mice (**Fig. 3b**). SH-273 eliminated STING-induced expansion of IL35^+^ and IL10^+^ Breg cells in the splenocytes from KPC mice (**Fig. 3c**). As a control, SH-273 did not show such effect in the splenic B cells from STING knockout mice (Tmem173^-/-^), suggesting SH-273 effect is indeed STING dependent (**Fig. S14**). In contrast, SH-273 sustained activity to stimulate IRF3 phosphorylation in myeloid cells, and thus SH-273 preserved STING function to activate these myeloid cells (**Fig. S12**)

**Fig. 3.**
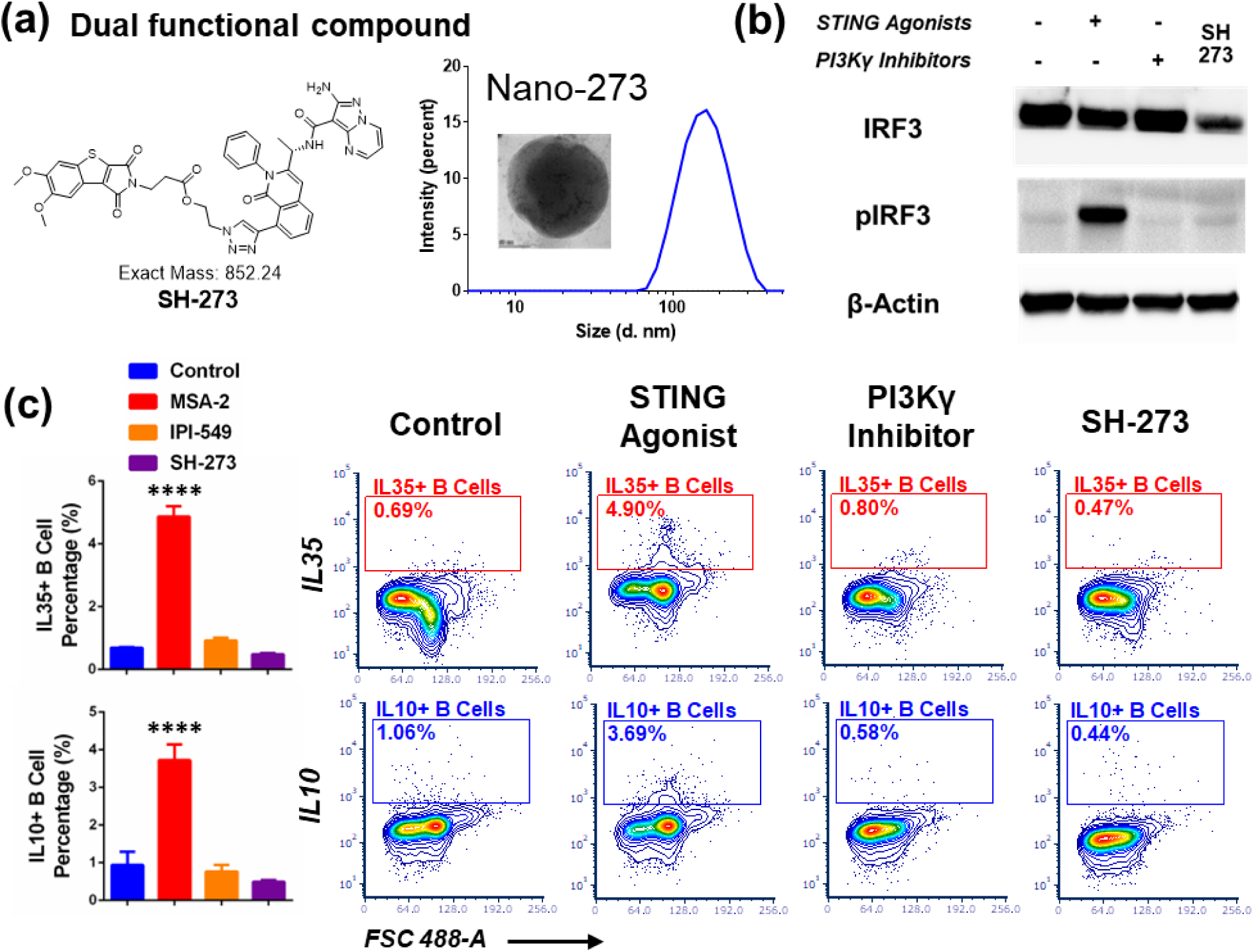
Dual functional compound SH-273 and its albumin nanoformulation eliminate Breg cells through abolishing STING-induced IRF3 phosphorylation. (**a**) Molecular structure of dual functional compound SH-273. Dynamic Light Scattering (DLS) size distribution of SH-273 albumin nano formulation (Nano-273). Insert image is Transmission electron microscopy (TEM) imaging of Nano-273. (**b**) Phosphorylation of IRF3 at splenic B cells derived from KPC PDAC mice after treatments with MSA-2, IPI-549 or SH-273, respectively. (**c**) Quantification and representative flow cytometry analysis of IL35^+^ and IL10^+^ Breg cells (from KPC spleen) after treatments with MSA-2, IPI-549 or SH-273, respectively. Data for quantification are shown as mean ± SD, n = 3. Statistical comparisons are based on one-way ANOVA, followed by post hoc Tukey’s pairwise comparisons or by Student’s unpaired T-test. The asterisks denote statistical significance at the level of **** p < 0.0001. ANOVA, analysis of variance; SD, standard deviation. For (**c**) statistical comparisons are conducted between MSA-2 group with other groups.

In order to achieve systemic immunity, we prepared an albumin nanoformulation of SH-273 (Nano-273) for intravenous administration and effective delivery to tumor and lymph nodes^11^. Nano-273 showed a uniform morphology with an average hydrodynamic diameter of 135 nm, a polydispersity index (PDI) of 0.095, maintaining stable even after 1000-fold dilution (**Fig. 3a and S15a**). Nano-273 showed effective delivery to pancreatic tumors and lymph nodes compared to SH-273 in transgenic KPC PDAC mice (**Fig. S15b**). Additionally, 2D and 3D confocal imaging showed that albumin nanoformulation of fluorescent dye delivered more and penetrate deeper into tumor tissue and tumor organoids from KPC PDAC mice than free fluorescent dye (**Fig S16-S17, Movie 1-3**). Furthermore, 2D confocal imaging revealed albumin nanoformulation of fluorescent dye delivered more and deeper into lymph nodes of KPC PDAC mice than free fluorescent dye (**Fig. S18)**. 3D confocal imaging showed albumin nanoformulation of fluorescent dye penetrated deep into the lymph nodes with accumulation in the macrophages and B cells (**Movie S4 – S9**).

### Nano-273 extended median survival to 200 days in KPC PDAC mice through activating systemic immunity

To assess the in vivo efficacy of Nano-273, we tested Nano-273 in combination with anti-PD1 in transgenic KPC PDAC mice with spontaneous pancreatic tumors. The data showed that the combination of Nano-273 and anti-PD-1 extended median survival to 200 days compared to control group (median survival at 120 days). MSA-2 combined with anti-PD-1 slightly extended median survival to 146 days. Neither the anti-PD-1 antibody alone nor combined with the albumin nanoformulation of paclitaxel (Nano-P) significantly improved median survival of KPC PDAC mice. (**Fig. 4a).** Nano-273 combined with anti-PD-1 showed superior efficacy in KPC mice compared to other treatment groups **(Fig S19**), which includes combination of IPI549 and MSA2, Nano-P and IPI549. At the endpoint of the study, we also examined the metastasis in the lung, liver, spleen, and kidney tissues. Notably, Nano-273 substantially decreased tumor metastasis and local invasion to lung (**Fig. 4b**).

**Fig. 4.**
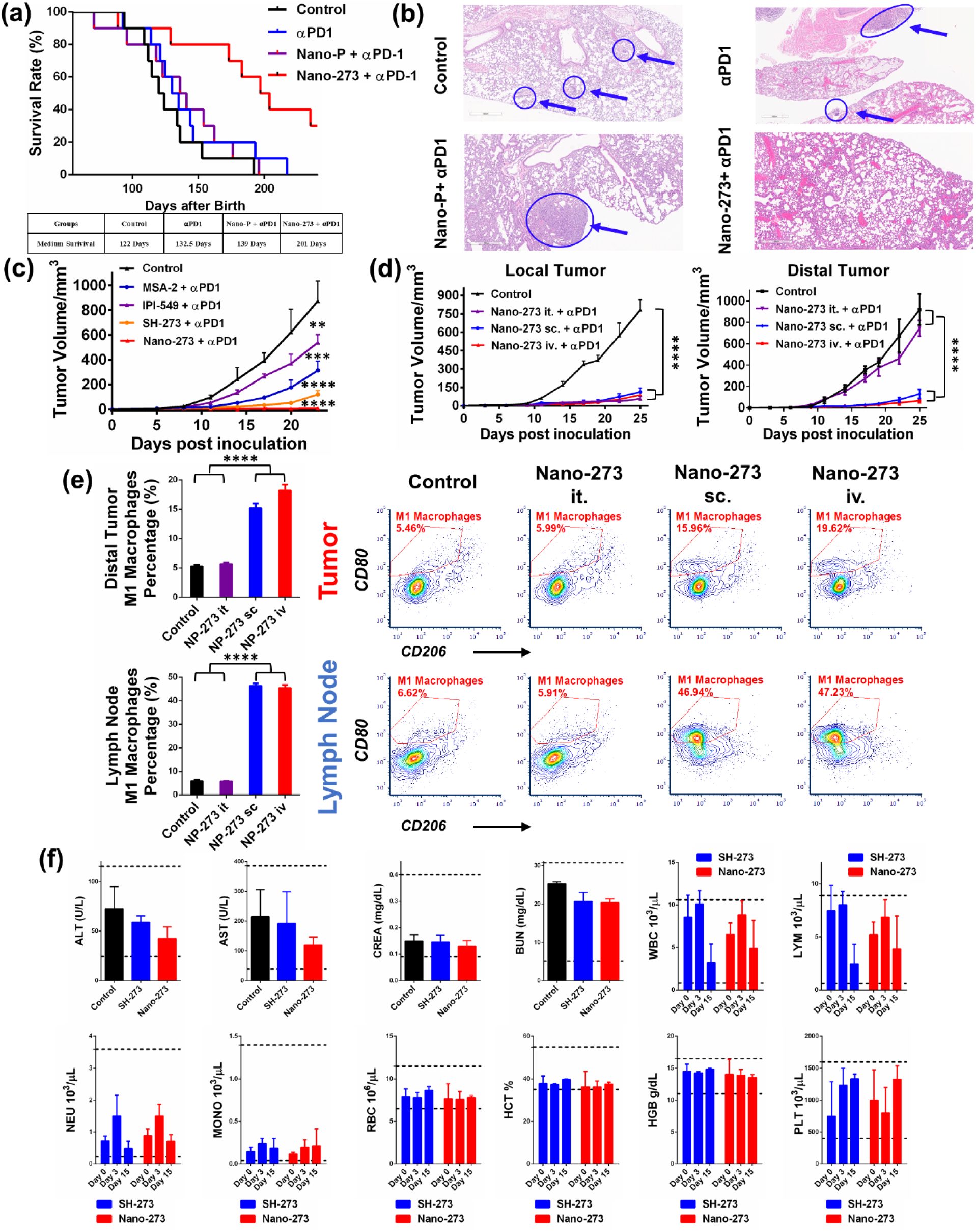
Nano-273 extended median survival to 200 days in KPC PDAC mice via activating systemic immunity without exhibiting toxicity. (**a**) Antitumor efficacy in transgenic KPC (LSL-KrasG12D, LSL-Trp53R172H/+, Pdx1cre/+) PDAC mice with different treatments. Survival rate of KPC mice after treatment with PBS (control), anti-PD-1 antibody (100 µg), Paclitaxel in albumin nano formulation (intravenous injection, 11.7 μmol/kg) plus anti-PD-1 antibody (100 µg) and Nano-273 (i.v. 17.6 μmol/kg) plus anti-PD-1 antibody (100 µg). n = 10 for each group. (**b**) Pathological staining of lung tissues from (**a**) to evaluate pancreatic cancer metastasis or invasion. Blue arrows and circles indicated metastasis or invasion foci. (**c**) Antitumor efficacy in C57BL/6 mice inoculated with KPC 6422 tumor after different treatments. Average tumor volume after treatment with PBS (control), MSA-2 (i.v. 34.0 μmol/kg) plus anti-PD-1 antibody (100 µg), IPI-549 (i.v. 18.9 μmol/kg) plus anti-PD-1 antibody (100 µg), SH-273 (i.v. 17.6 μmol/kg) plus anti-PD-1 antibody (100 µg) and Nano-273 (i.v. 17.6 μmol/kg) plus anti-PD-1 antibody (100 µg). The data represents the mean ± SD; n = 5. (**d**) Antitumor efficacy in C57BL/6 mice inoculated with KPC 6422 tumor with Nano-273 treatment in different administration routes. Average tumor volume of KPC pancreatic cancer mice after treatment with PBS (control) and Nano-273 (17.6 μmol/kg) after intratumorally (i.t), subcutaneous (s.c.) and intravenous (i.v.) injection, respectively plus anti-PD-1 antibody (100 µg). Local tumor is referred to the tumor with intra-tumor injection or near s.c. injection site. Distal tumor is referred to no intra-tumor injection or far from s.c. injection site. The data represents the mean ± SD; n = 5. (**e**) Quantification and representative flow cytometry analysis of M1 macrophage ratios at distal tumor and lymph node after treatments from (**c**). The data represents the mean ± SD; n = 3. (**f**) Liver enzymes, kidney function, and blood cell counts in CD1 mice after treatments with SH-273 and Nano-273 every 3 days for 5 doses (17.6 μmol/kg) through intravenous injection. The data represents the mean ± SD; n = 3. Statistical comparisons are based on one-way ANOVA, followed by post hoc Tukey’s pairwise comparisons or by Student’s unpaired T-test. The asterisks denote statistical significance at the level of * p < 0.05, ** p < 0.01, *** p < 0.001, **** p < 0.0001. ANOVA, analysis of variance; SD, standard deviation. For (**c**) statistical comparisons are conducted between treatment groups with control. Statistical comparisons in (**f**) showed no statistical significance.

In addition, we also tested the Nano-273’s efficacy in xenograft model using KPC 6422 cells in C57BL/6 mice. Both Nano-273 and SH-273, in combination with anti-PD-1 antibody, significantly delayed tumor growth compared to the MSA-2, IPI-549, and control groups, with Nano-273 demonstrating superior efficacy over free SH-273 (**Fig. 4c and S20**).

In order to evaluate the systemic anticancer immunity of Nano-273 through intravenous administration, we compared the anticancer efficacy of Nano-273 via intravenous (iv), subcutaneous (sc), or intratumorally (it) injections, to inhibit bilaterally inoculated KPC 6422 cells in C57BL/6 mice. Intratumorally injections of Nano-273 only suppressed local tumor growth, but had minimal inhibition to the tumor at the distal site. This phenomenon is confirmed in clinical studies that local intra-tumor injection of STING agonists only generated local tumor inhibition but did not have much effects on distal tumors or metastasis.^13,14,16–18^ This highlights the importance of using STING agonists systematically to induce systemic immunity. Indeed, our data showed that both intravenous and subcutaneous injections of Nano-273 elicited robust systemic immune responses, significantly reducing tumor burden at both local and distal sites (**Fig. 4d and S21**).

To investigate how systemic delivery of Nano-273 achieve systemic immunity in comparison with intratumoral injection, we used flow cytometry to analyze the M1-macrophage polarization in local, distal tumors and lymph nodes. The data showed that Nano-273 (iv and sc) significant increased M1 macrophages at distal tumors and lymph nodes, while intra-tumoral injection of Nano-273 did not increase M1-macrophage in either distal tumors or lymph nodes (**Fig. 4e**). To confirm if systemic efficacy of Nano-273 indeed depends on the systemic immune cell trafficking, we used fingolimod (FTY-720) to inhibit lymphocyte egress from lymph nodes^19^, which reversed effect of Nano-273 after iv administration (**Fig. S22**). Fingolimod also decreased immune infiltration in distal tumor and reduced M1 macrophage in the lymph nodes and distal tumor sites (**Fig. S23**). These data suggest that systemic delivery of Nano-273 achieve systemic immunity for its superior systemic anticancer efficacy in KPC PDAC mice.

Finally, to confirm if systemic delivery of Nano-273 will not cause toxicity in major organs, we performed extensive toxicity study using the same dose regimen as seen in the therapeutic studies (iv injection five doses of Nano-273 and SH273). No toxicity in the liver, kidney, and blood cells was observed as shown by liver enzymes (AST, GST), kidney function (BUN, creatinine) and blood cell counts (**Fig. 4f**). Pathological staining of major organs (liver, kidney, spleen, lung, and heart) also did not show toxicity as indicated healthy tissue structures (**Fig. S24**).

### Nano-273 eliminated STING-induced Breg cells expansion and remodeled immune microenvironment in tumor and lymph nodes for systemic anticancer immunity

To investigate how did Nano-273 remodel the tumor immune microenvironment, we first utilized flow cytometry to examine immune cell infiltration following the treatments in **Fig. 4a**. Both Nano-273 and SH-273 significantly increased immune cell infiltration to tumors compared to STING agonist (MSA-2) or PI3Kγ inhibitor (IPI-549) alone (**Fig. 5a**). Subsequently, we employed single-cell RNA-sequencing (RNA-Seq) to analyze all immune cells in the tumor tissues post-treatments. Immune cell phenotypes were identified based on RNA expression levels and corroborated by TotalSeq™-C antibody surface staining (**Fig. 5b and 5d**), which showed increased B and CD8 T cell populations. The data showed that both Nano-273 and SH-273 significantly decreased IL35^+^ and IL10^+^ Breg cells, while increased ICOSL+ B cell populations in tumors (**Fig. 5c, 5e and S25**)^20^. In contrast, STING agonist (MSA-2) significantly increased IL35^+^ and IL10^+^ Breg cells in tumors (**Fig. 5c, 5e and S25**), similar to literature reports^8,11,14^.

**Fig. 5.**
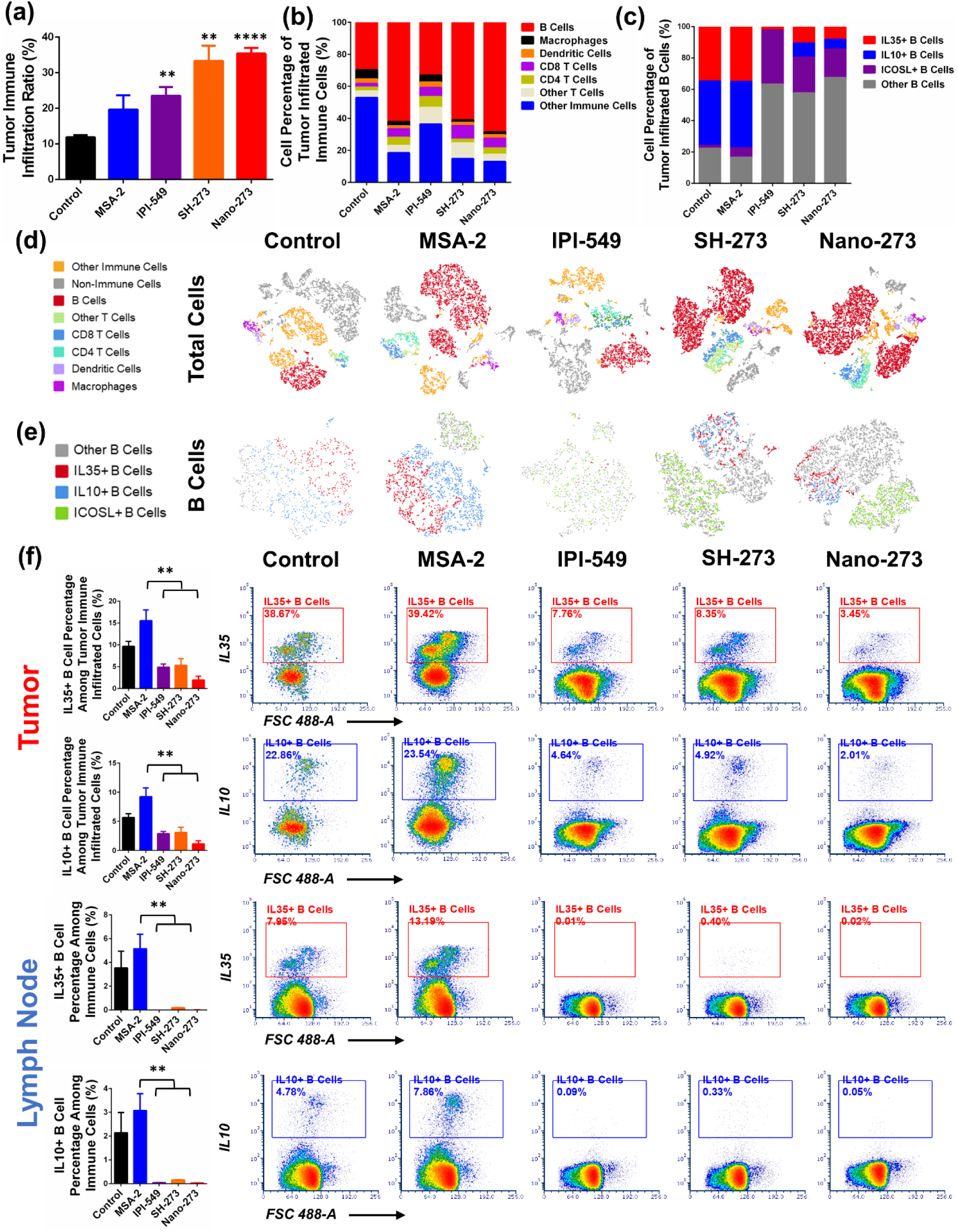
Nano-273 eliminated STING-induced Breg cells expansion and remodeled the immune microenvironment in tumor and lymph nodes for systemic anticancer immunity. (**a**) Quantification of tumor immune infiltration by flow cytometry from the residual tumors of the mice at 10 days after final dose in Fig. 4c. (**b**) Different immune cell populations among tumor infiltrating immune cells by single cell RNA-seq analysis from anti-tumor efficacy study in mice at 9 days after final dose in Fig. 4c. (**c**) Percentage of different subpopulations of B cells among total B cells by single cell RNA-seq analysis. (**d, e**) t-distributed Stochastic Neighbor Embedding (t-SNE) plots of tumor-infiltrating immune cells and B cells by single cell RNA-seq analysis from mice in anti-tumor efficacy study at Fig. 4c. (**f**) Quantification and representative flow cytometry analysis of IL35^+^ and IL10^+^ Breg cells at tumor and lymph node in mice from anti-tumor efficacy study at Fig. 4c. Statistical comparisons are based on one-way ANOVA, followed by post hoc Tukey’s pairwise comparisons or by Student’s unpaired T-test. The asterisks denote statistical significance at the level of ** p < 0.01, **** p < 0.0001. ANOVA, analysis of variance; SD, standard deviation. For (**a**) statistical comparisons are conducted between treatment groups with control.

To verify these findings in single cell RNA-seq, we further used flow cytometry to assess Breg cell frequencies in tumors, lymph nodes, and spleens post-treatment. The Nano-273 and SH-273 significantly decreased Breg cell frequencies by 1.8 to 7.8 -folds in tumors (IL35^+^ Breg cell: 2.0% and 5.3%, IL10^+^ Breg cell 1.2% and 3.1%), 10.5 to 510 - folds in lymph nodes (IL35^+^ Breg cell: 0.01% and 0.2%; IL10^+^ Breg cell: 0.03% and 0.2%) in comparison with MSA-2 and control groups in tumors (IL35^+^ Breg cell: 15.6% and 9.6%; IL10^+^ Breg cell: 9.3% and 5.7%), and in lymph nodes (IL35^+^ Breg cell: 5.1% and 3.5%; IL10^+^ Breg cell: 3.1% and 2.1%)(**Fig. 5f**). Our results suggest that Nano-273 effectively eliminated STING-induced Breg cells expansion and remodeled immune microenvironment in both tumors and lymph nodes, thereby achieving systemic anticancer immunity.

## Discussion

The immunosuppressive microenvironment in both tumor and lymph nodes of PDAC presents a significant hurdle for the efficacy of immunotherapy.^6,7^ Therefore, a variety of immune modulators targeting myeloid cell suppression have been developed^21–27^, but they have shown only limited clinical success^16,28^. Recent research showed that the immune suppression in PDAC’s is significantly regulated by high number of regulatory B cells in both tumors and lymphatic systems^8,9,11^. A well-known immune modulator, STING agonist, could activates myeloid cells, but unfortunately expands Breg cells in tumors and lymph nodes, contributing to therapeutic resistance^7,11,13,21–24^.

In this study, we discovered that the PI3Kγ inhibition during STING activation abolished IRF3 phosphorylation in B cells, effectively eliminated Breg cells, but PI3Kγ inhibition sustained IRF3 phosphorylation and thereby preserved STING activation in myeloid cells. Based on this finding, we developed a dual functional compound SH-273 to activate STING and inhibit PI3Kγ. SH-273 preserved STING function to activate myeloid cells and eliminated Breg cells to overcome STING resistance. To achieve systemic antitumor immunity, we also prepared an albumin nanoformulation Nano-273 for safe intravenous administration and effective delivery to pancreatic tumors and lymph nodes in transgenic KPC PDAC mice. Impressively, when combined with anti-PD-1 therapy, Nano-273 extended the median survival to 200 days in transgenic KPC PDAC mice. In contrast, other clinically used treatments did not significantly improve survival, including the anti-PD1 antibody alone, anti-PD1 combined with albumin nanoformulation of paclitaxel (nab-paclitaxel), and STING agonist MSA-2 with anti-PD1. Subsequent immune profiling using single cell RNA-seq and flow cytometry confirmed that Nano-273 markedly reduced Breg cell frequency and significantly remodeled the immune microenvironment than other treatment groups.

This study provides a novel and more effective strategy to overcome immune suppressive environment in both tumors and lymph nodes in PDAC tumors by simultaneously targeting myeloid cells and Breg cells. While the role of myeloid cells in immune suppression of PDAC is well-established, the significant impact of Breg cells has only recently been recognized^8–11^. Bhalchandra et al. found an increased Breg cells in primary human PDAC tissues from 56 patients compared to adjacent normal tissues^8^. Their findings revealed a negative relationship between Breg cells abundance in tumors and PDAC prognosis, but positive correlation between plasma cell numbers and PDAC prognosis. Similarly, Li et al. reported that STING agonist significantly increased the frequency of Breg cells in the pancreatic cancer mouse model^11^. Our study corroborates these findings, showing elevated Breg cells in PDAC mouse tumors relative to normal pancreatic tissue (**Fig. 1 and 5**), and we also confirmed that STING agonists increased Breg cells in PDAC (**Fig. 2 and 5**). This underscores the role of Breg cells in PDAC immune suppression and intrinsic resistance to STING agonists. Crucially, our research demonstrates that blocking PI3Kγ during STING activation eliminated STING-induced Breg cell expansion and significantly enhanced the effectiveness of immunotherapy in PDAC. Our data shows that Nano-273 markedly reduced IL35^+^ and IL10^+^ Breg frequency in tumor to less than 2% and 1.2%, compared to 15.6% and 9.3% in the group treated with a STING agonist, and 9.6% and 5.7% in the control group. Furthermore, Nano-273 triggered a more robust activation of the tumor immune microenvironment, evidenced by increased immune cell infiltration into tumors, and heightened frequencies of CD8 T cells, and M1 macrophages, compared to treatments with STING agonist, PI3Kγ inhibitor, or the control group.

While previous studies have emphasized the critical role of Breg cells in PDAC immune suppression, the specific signaling pathway to overcome Breg cell expansion during STING activation was not well-defined. Our research reveals that inhibiting PI3Kγ decreased IRF3 phosphorylation in B cells, hindering the expression of IL-35 and IL-10, thereby eliminating Breg cells, while it sustained IRF3 phosphorylation in myeloid cells, thus preserved STING activation for Type-I interferon production (**Fig. 2**). This suggests the divergent effects of PI3K during STING activation in Breg and myeloid cells^8,11^. PI3Kγ is mainly recognized for its role in regulating macrophage repolarization from M2 to M1 phenotype.^29,30^ PI3Kγ has also been reported to be expressed in B cells^31–34^. However, its function in B cells has been less understood.^34,35^ Our study presents a novel mechanism and potential therapeutic application for PI3Kγ inhibition in regulating Breg cells as a strategy in pancreatic cancer immunotherapy.

Furthermore, clinical trials using local intra-tumoral injections of STING agonists may only overcome immune suppression in the tumor microenvironment to achieve local anticancer immunity, but may not adequately address the lymph node microenvironment to achieve systemic anticancer immunity^13,14,16–18^. For instance, the local injection of STING agonist mainly shows anticancer efficacy in local tumors but does not exhibit strong anticancer efficacy in distal tumors or metastasis. Additionally, intra-tumoral injections are not feasible for many cancers in clinical setting, including pancreatic cancer.

Therefore, developing an effective systemic delivery for STING agonists has become a critical objective to achieve systemic anticancer immunity to improve their clinical efficacy^17,36,37^. In our study, we utilized albumin nanoformulation of SH-273 for safe intravenous administration and effective delivery to pancreatic tumors and lymph nodes. Clinically, albumin nanoformulation of paclitaxel (Abraxane^®^) has been approved as the first-line therapy in pancreatic cancer patients benefited from its superior efficacy compared to traditional solvent-based Taxol^®38^. Our previous research has also shown that albumin nanoformulations of immune modulators enhance the drug delivery to tumors and lymph nodes, thereby overcoming immune suppression in both tumors and lymphatic system^11^. Our findings show that intravenous administration of Nano-273 displayed no toxicity and effectively combatted both local and distal tumors (**Fig. 4**). Nano-273 exhibited greater accumulation in both tumors and lymph nodes compared to the solvent-based formulation (**Fig. S15b, S16-S18, Movie S1-S9**).

In conclusion, our dual functional compound SH-273, by stimulating STING and inhibiting PI3Kγ, preserves STING function in myeloid cells and eliminates Breg cells to overcome STING resistance. Nano-273 achieved systemic antitumor immunity through intravenous administration, which decreases Breg cells and remodels microenvironment in tumors and lymph nodes. Nano-273, combined with anti-PD-1, extended median survival of 200 days in transgenic KPC PDAC mice (KrasG12D-P53R172H-Cre). Nano-273 is a promising candidate for treatment of PDAC.

## Methods

### Animal experiments

All animal experiments were conducted according to protocols approved by the University of Michigan Committee on Use and Care of Animals (UCUCA). Animals were maintained under pathogen-free conditions, in temperature- and humidity-controlled housing, with free access to food and water, under a 12-hour light-dark rhythm at the unit for laboratory animal medicine (ULAM), part of the University of Michigan medical school office of research. CD1 and C57BL/6 mice were obtained from The Jackson Laboratories. The KPC (LSL-KrasG12D/+;LSL-Trp53R172H/+;Pdx-1-Cre) transgenic mice were bred, genotyped and maintained by ULAM following reference^39^. Tumor experiments by xenograft implanting were performed using 6-week-old male wild-type C57BL/6 mice. Endpoints for anti-tumor efficacy studies were determined using the End-Stage Illness Scoring System, mice receiving an End-Stage Illness Score greater than 6 were euthanized by CO2 asphyxiation.

### Cells

All cells were maintained at 37 °C in a 5% CO2/95% air atmosphere and 85% relative humidity. Primary B-cells, CD4 T cells and splenocytes were cultured in RPMI-1640 media supplemented with 10% fetal bovine serum, 2-Mercaptoethanol (50 µM) and 1% pen/strep. THP-1-Blue™ ISG, THP-1 hSTING^KO^ and THP-1 hSTING^R232^ cells were cultured in RPMI 1640 supplemented with 2 mM L-glutamine, 25 mM HEPES, 10% heat-inactivated fetal bovine serum, 100 µg/mL normocin and 1% pen/strep. Bone marrow-derived dendritic cells (BMDCs) and bone marrow-derived macrophages (BMDMs) were isolated from mouse bone marrow cells and cultured following previously reported protocols^40,41^. RAW264.7 macrophages were cultured in complete RPMI-1640 media supplemented with 10% fetal bovine serum, 1% L-glutamine, 1% MEM nonessential amino acid solution, 1% sodium pyruvate and 1% pen/strep. KPC 6422 cells obtained from kerafast were cultured in DMEM supplemented with 10% fetal bovine serum, Glutamax and 1% pen/strep^42^.

### Analyze Breg cells and macrophages in various tissues and assess the effectiveness of STING agonist in pancreatic cancer mouse models

To analyze Breg cells and macrophage proportions, lymph node, tumor, and spleen tissues from KPC (LSL-Kras G12D, LSL-P53R172H, Pdx1-cre) transgenic mice were harvested and prepared for single-cell suspension. CD45, CD4, CD19, IL35, and IL10 were used to identify regulatory B cells by flow cytometry. Similarly, CD45, CD11b, F4/80, CD80, and CD206 were used to identify M1 and M2 macrophages. To evaluate the in vivo efficacy of the STING agonists, 6-week-old C57BL/6 male mice were subcutaneously inoculated at the right flank with 5*10^5 KPC 6422 cells/mouse. DiABZi (20 μg/mouse for intratumorally injection and 1.5 mg/kg for intravenous injection) and 2’3’-cGAMP (10 μg/mouse for intratumorally injection) were administrated 5 days post tumor inoculation, every 3 days for a total of 5 times. Tumor volumes were calculated as volume = (width)^2 × length/2.

### Western blot analysis of STING and PI3Kγ

Lymph node, pancreas and tumor were harvest from C57BL/6 mice or transgenic KPC mice. Proteins were immediately extracted post-harvest for western blot analysis. B cells were isolated from splenocytes of C57BL/6 mice or transgenic KPC mice using the STEMCELL EasySep™ mouse B cell isolation kit. Bone marrow-derived dendritic cells (BMDCs) and bone marrow-derived macrophages (BMDMs) were isolated from bone marrow cells of C57BL/6 mice or transgenic KPC mice and cultured following previously reported protocols^40,41^. Proteins were prepared following cell lysis, sonication, and centrifugation for western blot analysis.

### Study the role of PI3Kγ in STING activation in different immune cells

B and CD4+ T cells were isolated from the spleen using the STEMCELL EasySep™ mouse B cell isolation kit and EasySep™ mouse CD4+ T cell isolation kit, respectively. To investigate the phosphorylation of IRF3, B cells (pre-treated with 5 µg/mL anti-IgM), CD4+ T cells, BMDCs, BMDMs, and THP-1-Blue™ ISG cells were seeded in 12-well plates at a density of 3*10^6 cells/well and incubated with or without MSA-2 (5 µg/mL) and with or without IPI-549 (17 µM). Proteins were analyzed by western blot analysis after cell collection and lysis. IL35+ or IL10+ cells were analyzed by flow cytometry after cell collection and staining with anti-EBI3 and anti-IL10 antibodies. For STING activation in BMDCs, BMDMs, THP-1 cell lines, and RAW264.7 cells, the cells were treated with or without 10 µM of MSA-2 and with or without 10 µM of IPI-549. Supernatants were collected for cytokine measurement via ELISA. BMDCs were collected and stained with CD80 and CD86 for activation analysis by flow cytometry. BMDMs and RAW264.7 cells were collected and stained with CD80 or CD86 to access M1 polarization ratios by flow cytometry.

### Synthesis of SH-273, preparation of albumin nanoformulation of SH-273, and test of their cellular and binding activity

Synthesis of SH-273 is described in **Scheme 1**. Starting material compound **1** was cyclized with methyl thioglycolate to afford **2**. Then, condensation between **2** and amine gave **3** which was brominated with NBS to produce **4**. Sequential oxidation of **4** by NMO and PCC produced **6** followed by deprotection to furnish intermediate **7.**^43^

Starting material compound **8**, (S)-2-amino-N-(1-(8-ethynyl-1-oxo-2-phenyl-1,2-dihydroisoquinolin-3-yl)ethyl)pyrazolo[1,5-a]pyrimidine-3-carboxamide reacted with 2-azidoethanol in CH_2_Cl_2_ with CuI.A-21 catalyst to afford intermediate **9**. The reaction mixture was sonicated for 1h and stirred at room temperature overnight. Upon completion of the reaction, the catalyst was removed by filtration using CH_2_Cl_2_ and the solvent was evaporated under vacuum.

The solution of intermediate **9** (66.9 mg, 0.125 mmol), **7** (44 mg, 0.13 mmol), HATU (71.3 mg, 0.187 mmol), DIPEA (24.2 mg, 0.187 mmol), and DMAP (15 mg, 0.125 mmol) in DMF (1 mL) was stirred at room temperature for 38 h. Upon completion of the reaction, the mixture was concentrated to yield SH-273, which was then purified using column chromatography (20:1 ethyl acetate/methanol) to provide the title compound **SH-273** (74.55 mg).

NMR of SH-273: ^1^H NMR (599 MHz, CDCl_3_) δ 8.40 (dd, *J* = 6.7, 1.7 Hz, 1H), 8.36 (dd, *J* = 4.5, 1.6 Hz, 1H), 8.12 (s, 1H), 7.95 (s, 1H), 7.88 (d, *J* = 7.0 Hz, 1H), 7.78 (dd, *J* = 7.6, 1.4 Hz, 1H), 7.55 (t, *J* = 7.7 Hz, 1H), 7.43 (d, *J* = 8.9 Hz, 3H), 7.39 – 7.33 (m, 3H), 7.23 (s, 2H), 6.75 (dd, *J* = 6.8, 4.5 Hz, 1H), 6.62 (s, 1H), 5.53 (s, 2H), 4.72 (t, *J* = 6.9 Hz, 1H), 4.57 – 4.49 (m, 2H), 4.43 (t, *J* = 5.2 Hz, 2H), 3.92 (s, 6H), 3.73 – 3.67 (m, 1H), 3.68 – 3.60 (m, 2H), 3.11 (dd, *J* = 7.5, 4.3 Hz, 1H), 2.73 (s, 3H), 2.48 (t, *J* = 6.5 Hz, 2H), 1.40 (dd, *J* = 6.9, 5.0 Hz, 4H), 1.35 (dd, *J* = 9.4, 6.7 Hz, 4H).

Albumin nano formulation of SH-273 (Nano-273) was then prepared followed previously established protocol^11^. Briefly, SH-273 (10 - 12 mg) was dissolved in 1mL of chloroform and then mixed with mouse albumin solution (100 mg/20 mL) to generate a milky emulsion using a rotor-stator homogenizer. This emulsion was processed through five to six cycles at 26,000 psi in a high-pressure homogenizer (Nano DeBEE) at 4°C. Subsequently, the organic solvent was removed using a rotary evaporator at 25°C. The Nano-273 suspension was then filtered (0.22 μm), lyophilized, and stored at −20°C. Size distribution and morphology were assessed using dynamic light scattering and transmission electron microscopy (TEM).

The cellular IC50 of SH-273 was measured using THP-1-Blue™ ISG cells. THP-1-Blue™ ISG cells were seeded in a 96-well plate with a density of 1*10^5 cells/well and incubated with different concentrations of MSA-2 and Nano-273 for 24 hrs at 37 °C. Cell media were then collected, mixed, and incubated with QUANTI-Blue™ solution. SEAP levels were subsequently measured using Synergy 2 microplate reader (Biotek) at 620 nm in absorbance mode. B cells (pre-treated with 5 µg/mL anti-IgM) and Bone marrow dendritic cells were seeded in a 12-well plate at a density of 3*10^6 cells/well and incubated with SH-273 (17 µM). Proteins were analyzed by western blotting after cell collection and lysis. For IL35 or IL10 expression analysis, cells were collected and stained with anti-EBI3 and anti-IL10 antibodies for flow cytometry.

PI3Kγ binding affinity was evaluated using PI3Kγ (p110γ/PIK3R5) assay kit from BPS Bioscience, following the provided protocol. Inhibitors were prepared at 5X concentration in an aqueous solution at various concentrations. The PI3Kγ enzyme was diluted to ∼4 ng/μl with 2.5x Kinase assay buffer. Reaction components were added sequentially: 5 μl of PI3K lipid substrate, 5 μl of inhibitor or inhibitor buffer, 5 μl of 12.5 μM ATP, and 10 μl of diluted PI3Kγ. Between each addition, the plate was shaken for 1 minute to ensure thorough mixing. After a 40-minute reaction, 25 μl of ADP-Glo reagent was added to each well. The plate was covered with aluminum foil and incubated at room temperature for 45 minutes. Subsequently, 50 μl of Kinase Detection reagent was added to each well, followed by another 30-minute incubation at room temperature, again covered with aluminum foil. Luminescence was measured using a microplate reader.

### Pharmacokinetics and toxicity of SH-273 and Nano-273 in mice

To investigate the pharmacokinetics of SH-273 and Nano-273, KPC transgenic mice (approximately 100 days old with spontaneous tumors) were divided into two groups (n = 9) and treated with SH-273 (i.v. 17.6 μmol/kg) or Nano-273 (i.v. 17.6 μmol/kg). Tissue samples (plasma, liver, tumor, lymph nodes) were collected at 0.5, 2, and 7 hours post-administration, processed, and analyzed. SH-273 concentrations were measured by LC-MS using An AB SCIEX QTRAP 5500 mass spectrometer with a Turbo V electrospray ionization source (Applied Biosystems) coupled to a Shimadzu LC-20AD HPLC system, was used for the quantification. HPLC separation was performed on a waters XBridge® C18 3.5 μm, 5 cm x 2.1 mm column (Waters Corporation). Mobile phase A (water containing 0.1% formic acid) was first kept at 95% for 0.8 min, then decreased to 1% during 1.2 min, and maintained for 1.5 min, returned to 5% B (acetonitrile containing 0.1% formic acid) and maintained for 2 min. The flow rate was 400 µL/min. Positive ion MS/MS was conducted to detect SH273. The mass spectrometric conditions were set as follows: source temperature, 500°C; curtain gas (CUR), 30 psi; ion spray voltage, 5500 V; ion source gas 1 (GS1), 60 psi; ion source gas 2 (GS2), 40 psi; collision gas (CAD), high; entrance potential (EP), 10 eV; collision energy (CE), 45 eV. The MS/MS transition 853.2 > 648.1 was used to detect SH273, and a dwell time of 50 ms was used for the transition. And the second transition 473 > 431.1 was used for the internal standard Clofazimine (CFZ). Data acquisition and quantitation were performed using Analyst software (Applied Biosystems). To determine the SH273 in mice plasma, 140 μL of internal standard solution (CFZ 50 ng/mL in Acetonitrile) and 30 μL of acetonitrile were added into 30 μL of plasma samples. The mixture was vortexed for 10 min and centrifuged at 3500 rpm for 10 min.

Then, the supernatant was transferred to the autosampler vials for LC–MS/MS analysis. tissue samples were homogenized (Precellys tissue homogenizer, Bertin Technologies) with the addition of 20% Acetonitrile-water solution with a ratio of 5:1 volume (mL) to weight of tissue (g). Then, 140 μL of internal standard solution and 30 μL of acetonitrile were added into 30 μL of tissue homogenization samples for protein precipitation. The mixture was vortexed for 10 min and centrifuged at 3500 rpm for 10 min. The supernatant was transferred to the autosampler vials for LC–MS/MS analysis. Blank plasma and tissues, the samples from un-treated control groups, were used to exclude contamination or interference. The SH273 analytical curves were constructed with 12 standards spiked with blank plasma or tissue, by plotting the peak area ratio of SH273 to the internal standard versus the sample concentration. The concentration range evaluated was from 1 to 5000 ng/mL in plasma and tissues. All samples for calibration curve were made as the method mentioned above.

To study the in vivo toxicity, CD1 mice (6 weeks) were randomly assigned to two groups (n = 4) and administered with SH-273 (15 mg/kg) or Nano-273 (15 mg/kg) every three days for 5 times. Blood was collected on day 0, day 3, and day 15 following the initial dose for whole blood cell count. At day 15, plasma was collected for test of liver enzymes and kidney functions. At day 15, organs (heart, liver, spleen, lung and kidney) were harvest and collected for pathological staining. All blood cell count, liver enzymes, kidney functions and tissues pathological staining were conducted by ULAM.

### In vivo anti-tumor efficacy

KPC transgenic mice were randomly assigned to various treatment groups (n = 10). Treatments started at 8 weeks of age, with doses administered weekly for a total of five times. The survival dates of the mice were documented either at the time of death or upon reaching the endpoint as defined by the End-Stage Illness Scoring System. Lung tissues were collected and underwent pathological staining to analyze metastasis in KPC mice that reached the endpoint in each treatment group by ULAM. C57BL/6 male mice at 6 weeks of age were subcutaneously inoculated with 5*10^5 KPC 6422 cells at the right flank (**Fig. 4c**) or both flanks (**Fig. 4d**). MSA-2 (i.v. 34.0 μmol/kg) plus anti-PD-1 antibody (100 µg), IPI-549 (i.v. 18.9 μmol/kg) plus anti-PD-1 antibody (100 µg), SH-273 (i.v. 17.6 μmol/kg) plus anti-PD-1 antibody (100 µg) and Nano-273 (i.v. 17.6 μmol/kg) plus anti-PD-1 antibody (100 µg) were then administered to the mice 5 days post-tumor inoculation and repeated every 3 days for a total of five times. Tumor volumes were calculated as volume = (width)^2 × length/2.

### Immune profiling of lymph nodes and tumors after treatments in vivo by flow cytometry

For in vivo M1 macrophage ratio measurements, lymph nodes and distal tumors were harvested and processed into single-cell suspensions 10 days following the final dosage. Markers CD45, CD11b, F4/80, CD80, and CD206 were used to identify M1 macrophages by flow cytometry. For in vivo Breg cell measurements, lymph nodes and tumors were harvested and processed into single-cell suspensions 10 days after the final dosage. CD45, CD4, CD19, IL35, and IL10 markers were used to identify regulatory B cells by flow cytometry.

### Single-cell RNA sequencing for immune fingerprints of tumors after treatments in vivo

Tumor samples were harvested and dissociated into single-cell suspensions 9 days after the last dose. Dead cells were excluded using a dead cell removal kit (Miltenyi Biotec). The single-cell suspensions were then stained with TotalSeq™-C Mouse antibody. These suspensions were subjected to final cell counting on a Countess II Automated Cell Counter (Thermo Fisher) and diluted to a concentration of 700-1000 nuclei/µL. We constructed 3′ single-nucleus libraries using the 10x Genomics Chromium Controller and followed the manufacturer’s protocol for 3′ V3.1 chemistry with NextGEM Chip G reagents (10x Genomics). The final library quality was assessed using a TapeStation 4200 (Agilent), and the libraries were quantified by Kapa qPCR (Roche). Pooled libraries were subjected to 150 bp paired-end sequencing according to the manufacturer’s protocol (Illumina NovaSeq 6000). Bcl2fastq2 Conversion Software (Illumina) was used to generate demultiplexed Fastq files, and a CellRanger Pipeline (10x Genomics) was used to align reads and generate count matrices.

### Immunofluorescence staining of tumor and lymph node

For immunofluorescence staining, tumor and lymph node tissues from transgenic KPC mice were harvested 5 hours post-intravenous administration of fluorescent dye (50 mg/kg). The tissues were immediately fixed in 1% paraformaldehyde for 1 hour, followed by immersion in 30% sucrose in 0.1% NaN3 in PBS overnight. Subsequently, tissues were embedded in optical coherence tomography (OCT) compound and snap-frozen in a CO2(s)+EtOH bath. Tissue sections of 15 µm were prepared and air-dried for 0.5 hours prior to staining. The sections were incubated with a blocking solution for 1 hour, followed by staining solution overnight at 4°C. Slides were then mounted with VECTASHIELD^®^ Mounting Medium (lymph node) and ProLong™ Diamond Antifade Mountant with DAPI (tumor) for confocal imaging. F4/80 (Alexa Fluor 594) was used to stain macrophages at tumor. B220 (Brilliant Violet 421), CD 3 (Alexa Fluor 488) and F4/80 (Alexa Fluor 647) were used to stain B cells, T cells and macrophages at lymph node, respectively.

### 3D imaging of in vivo lymph node and tumor distribution

Tumors and lymph nodes were collected 5 hours after intravenous administration of fluorescent dye (50 mg/kg). Harvested tissues were immediately processed following iDisco protocol for clearing and staining of antibodies. For tumor immune fluorescence staining, F4/80 (Alexa Fluor 594) and B220 (Brilliant Violet 421) were validated and used. For lymph node immune fluorescence staining, B220 (Brilliant Violet 421), F4/80 (Alexa Fluor 594) and CD 3 (Alexa Fluor 488) were validated and used. Subsequent 3D imaging of the tumors and lymph nodes was performed using Zeiss Light Sheet 7 and Nikon N-SIM Super resolution + A1R Confocal microscopes, respectively.

#### Sample pretreatment with methanol

1. Dehydrate with methanol/H2O series: 20%, 40%, 60%, 80%, 100%; 1h each.
2. Wash further with 100% methanol for 1h and then chill the sample at 4°c.
3. Overnight incubation, with shaking, in 66% DCM / 33% Methanol at room temperature.
4. Wash twice in 100% Methanol at room temperature, and then chill the sample at 4°C.
5. Bleach in chilled fresh 5%H2O2 in methanol (1 volume 30% H2O2 to 5 volumes MeOH), overnight at 4°C.
6. Rehydrate with methanol/H2O series: 80%, 60%, 40%, 20%, PBS; 1h each at room temperature.
7. Wash in PTx.2 1hr for two times at room temperature.

#### Immunolabeling

1. Incubate samples in permeabilization solution, 37°C, **n**/2 days (max. 2 days)
2. Block in blocking solution, 37 °C, **n**/2 days (max. 2 days).
3. Incubate with primary antibody in PTwH/5%DMSO/3% donkey serum, 37°C, **n** days.
4. Wash in PTwH for 4-5 times until the next day.

#### Clearing

1. Dehydrate in methanol/H2O series: 20%, 40%, 60%, 80%, 100%, 100%; 1 hour each at room temperature.
2. 3 hours incubation, with shaking, in 66% DCM / 33% methanol at room temperature.
3. Incubate in 100% DCM 15 minutes twice (with shaking) to wash the MeOH.
4. Incubate in dibenzyl ether (no shaking).

PTx.2 buffer (1 L): 100 mL PBS 10X, 2 mL TritonX-100.

PTwH buffer (1 L): 100 mL PBS 10X, 2 mL Tween-20, 1 mL Heparin stock solution (10 mg/mL).

Permeabilization solution (500 mL): 400 mL PTx.2 buffer, 11.5 g of glycine, 100 mL of DMSO.

Blocking solution (50 mL): 42 mL PTx.2 buffer, 3 mL of donkey serum, 5 mL of DMSO.

## Data availability

The data that support the plots in this paper and other findings of this study are available from the corresponding author upon reasonable request.

## Acknowledgments

We acknowledge the use of shared resource facility at the University of Michigan (Pharmacokinetics, Flow cytometry, Microscopy, Advanced Genomics, and In Vivo Animal cores).

